# Stimulus-Specific Information Flow Across the Canonical Cortical Microcircuit

**DOI:** 10.1101/753368

**Authors:** David A. Tovar, Jacob A. Westerberg, Michele A. Cox, Kacie Dougherty, Thomas Carlson, Mark T. Wallace, Alexander Maier

**Affiliations:** Neuroscience Program, Vanderbilt University, Nashville, TN, 37240, USA; School of Medicine, Vanderbilt University, Nashville, TN, 37240, USA; Department of Psychology, Vanderbilt University, Nashville, TN, 37240, USA; Center for Integrative and Cognitive Neuroscience, Vanderbilt University, Nashville, TN, 37240, USA; Vanderbilt Vision Research Center, Vanderbilt University, Nashville, TN, 37240, USA; School of Psychology, University of Sydney, Sydney, Australia; Department of Hearing and Speech Sciences, Vanderbilt University, Nashville, TN, 37240, USA; Department of Psychiatry, Vanderbilt University, Nashville, TN, 37240, USA; Kennedy Center for Research on Human Development, Vanderbilt University, Nashville, TN, 37240, USA

**Keywords:** Vision, Cortical Circuits, Machine Learning, Decoding, Multivariate Pattern Analysis

## Abstract

The vast majority of mammalian neocortex consists of a stereotypical microcircuit, the canonical cortical microcircuit (CCM), consisting of a granular input layer, positioned between superficial and deep layers. Due to this uniform layout, neuronal activation tends to follow a similar laminar sequence, with unique information extracted at each step. For example, the primate primary visual cortex (V1) combines the two eyes’ signals, extracts stimulus orientation and modulates its activity depending on stimulus history. Several theories have been proposed on when and where these processes happen within the CCM’s laminar activation sequence, but it has been methodologically challenging to test these hypotheses. Here, we use time-resolved multivariate pattern analysis (MVPA) to decode information regarding the eye-of-origin, stimulus orientation and stimulus repetition from simultaneously measured spiking responses across V1’s laminar microcircuit. We find that eye-of-origin information was decodable for the entire duration of stimulus presentation, but diminished in the deepest layers of V1, consistent with the notion that two eyes’ signals are combined within the upper layers. Conversely, orientation information was transient and equally pronounced across the microcircuit, in line with the idea that this information is relayed to other areas for further processing. Moreover, when stimuli were repeated, information regarding orientation was enhanced at the expense of eye-of origin information, suggesting that V1 modulates information flow to optimize specific stimulus dimensions. Taken together, these findings provide empirical evidence that adjudicates between long-standing hypotheses and reveals how information transfer within the CCM supports unique cortical functions.

**Significance Statement:** Despite the brain’s daunting complexity, there are common organizing principles across brain areas. For example, neocortical activation follows a stereotypical pattern that spreads from input layers towards layers above and below. While this activation pattern is well known, it has been challenging to ascertain how unique types of information are extracted within this common sequence in different brain areas. Here we use machine learning to track the flow of stimulus-specific information across the layers of visual cortex. We found that information regarding several separate stimulus dimensions was routed uniquely within the common activation sequence in a manner that confirmed prior model predictions. This finding demonstrates how differences in information flow within the stereotypical neocortical activation sequence shape area-specific functions.

## Introduction

Most cortical processing occurs within a stereotypical neuronal microcircuit termed the canonical cortical microcircuit (CCM; (1)). The CCM gives rise to a series of distinct, yet overlapping, activation steps that are spatially segregated between the superficial (supragranular), deep (infragranular) and middle (granular) layers of cortex (1–5). Ascending (feedforward) signals from parts of the brain that are closer to the sensory periphery terminate in the middle layers of cortical areas while descending (feedback) signals from downstream areas target the layers above and below (2, 3, 6).

Since the CCM is shared across most of the neocortex, an improved understanding of the laminar cortical processing chain is bound to translate into an improved understanding of cortical processing more generally (1, 6–9). However, our knowledge about laminar neuronal activation is rather limited (10). While recent studies demonstrated that there are two distinct sequences of laminar activation for feedforward and feedback activation, respectively (11–14), little is known about the different types of processes that occur along the CCM. For example, there are several open questions regarding how stimulus information is extracted along the primate primary visual cortex’s (V1) laminar microcircuit. Specifically, information regarding both stimulus orientation and the combination of the two eyes’ signals have been suggested to mainly occur in the superficial layers, one step removed from the initial point of activation in the middle layers (7, 15). However, the hypothesis that the two eye’s signals stay segregated until they arrive in the upper layers of V1 remains debated (16–20). In the same vein, several studies have postulated that orientation tuning originates before or during activation of the middle layers of V1 (21–26).

It has been technically challenging to directly test these competing hypotheses. Moreover, it is largely unknown how context such as stimulus history affects these processes. Using multi-electrode recordings, experimenters routinely assess neural activity across cortical layers simultaneously. The information amassed using these techniques far exceeds what can be gathered using single electrodes (27). However, laminar recordings are frequently analyzed using the same univariate techniques that have been established for single electrodes, rather than utilizing additional information from neighboring electrode contacts. This analytical shortcoming can be overcome by using statistical approaches that quantify information distributed across neighboring measurements in the brain, directly capturing neuronal interactions on the population level. Multivariate pattern classification analysis (MVPA) in particular has proven fruitful in systems neuroscience (28–31). More recently, time-resolved MVPA has emerged as a powerful technique to study the time courses with which information processing occurs across the brain (32, 33). While time-resolved MVPA has been applied to multielectrode recordings (34), to date no study to our knowledge has used this technique to analyze laminar cortical activation. Here we use time-resolved MVPA in primate primary visual cortex (V1) to test the competing hypotheses about the laminar flow of information outlined above.

## Results

We recorded population spiking responses across all layers of primary visual cortex (V1) of two macaque monkeys that passively viewed a randomized series of grating stimuli (Fig. 1). We repeated this experiment for 48 experimental sessions (n=8 for monkey E). For each of these sessions we penetrated the animals’ dura mater with a linear multielectrode array and ensured that it spanned all layers of V1 (see Methods and Suppl. Fig. 1). Extracellular voltages were measured every 100 microns at a 30 kHz sampling rate. Using current source density (CSD) and other physiological criteria (see Methods), we determined proper laminar electrode alignment and located the main retino-geniculate input layer (layer 4C in primates). Next, we mapped receptive fields using a reverse-correlation technique (see Methods and Suppl. Fig. 1A). Since achieving single-unit isolation on every channel is difficult, we instead opted to estimate the local population spiking response by quantifying the time-varying activity in the spiking frequency range (multi-unit activity, MUA).

**Figure 1:**
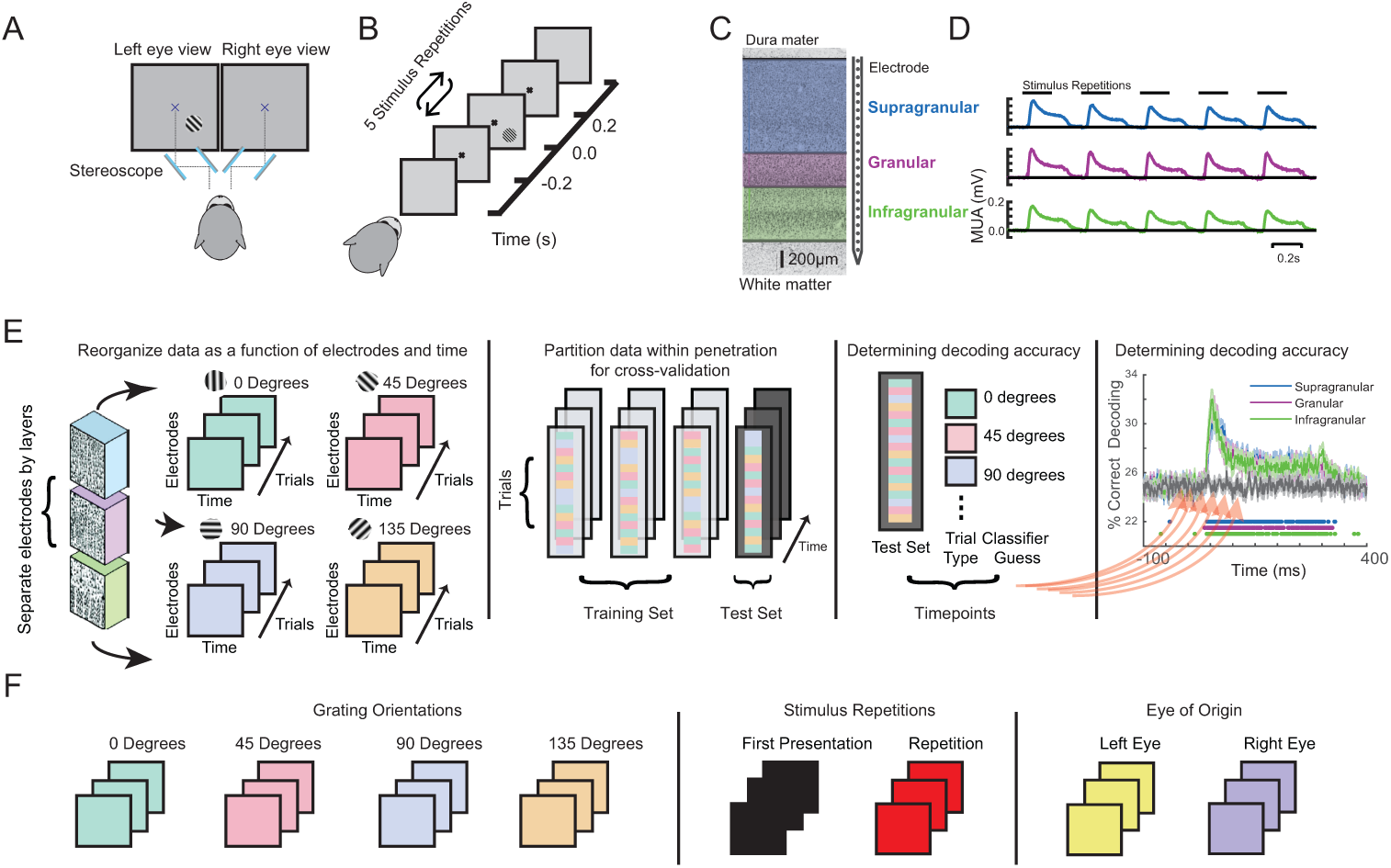
(A) Experimental setup. Monkeys were positioned in front of a monitor and tasked to passively fixate a central dot through a custom mirror stereoscope. (B) Experimental paradigm. Monkeys were shown a series of five grating stimuli of randomly varying orientations and ocular configuration with all other parameters were held constant. (C) Linear multicontact array recording laminar neuronal responses at 100 micron spatial resolution spanning through visual cortex. (D) Grand average multiunit spiking responses (MUA) to the stimulus sequence for all three main laminar compartments (both animals, all sessions). (E) Schematic of multivariate pattern analysis (MVPA). Population spiking responses (MUA) from each laminar compartment were reorganized as a function of electrode contact and time. A classifier was trained at each timepoint using linear discriminant analysis and 4-fold cross validation. (F) Decoded stimulus dimensions. Decoding analysis was separately performed for grating orientations, stimulus history (initial stimulus vs. repetitions), and eye-of-origin.

We placed the monkeys in front of a mirror-stereoscope to independently stimulate each eye (Fig.1A). We proceeded to present monocular visual stimuli to the neurons’ receptive field location while the animals kept fixation. Each stimulus consisted of a static sine-wave grating of fixed contrast, phase and spatial frequency, but randomly varying orientation and eye-of-origin (see Methods). For each trial, five of these gratings were presented for 200 ms each, interrupted by a 200 ms interstimulus interval (Fig., 1B; Suppl. Fig. 1B). Following each trial, the animals received liquid reward and were free to move their gaze. After a variable delay, animals selfinitiated the next trial by acquiring fixation.

Using neurophysiological criteria, we resolved the laminar MUA responses collected during this task into three main laminar compartments (see Methods; Fig. 1C). The granular compartment (layer 4C), shown in purple, receives the main retino-geniculate inputs from the eyes. Neurons in the granular compartment mainly project to the supragranular layers, shown in blue. Supragranular neurons predominantly target neurons in the infragranular compartment, shown in green (2). The resulting sequence of laminar activation following visual stimulation is captured by the canonical microcircuit model (CCM) (1, 4, 35, 36), which serves as the basis for a multitude of influential computational models of cortical function (6, 37–40). Figure 1D shows the grand average supragranular, granular and infragranular population spiking responses to the entire stimulus sequence (trial containing 5 stimulus presentations), across all recording sessions and both animals (41).

To track how sensory information from different stimulus dimensions are processed within this laminar microcircuit, we applied multivariate pattern analysis (MVPA) to the MUA of each of the three laminar compartments (Fig. 1E, left-most panel). To do so, we assembled two-dimensional neuronal response matrices (NRMs) that contained the millisecond-by-millisecond population spiking response at each electrode channel as a function of trials (see Methods). The specific stimulus dimensions we tested comprised of grating orientation, the eye that the stimuli were presented to (eye-of-origin) and the relative position of each stimulus within the stimulation sequence (Fig. 1F). We next randomly divided trials within sessions to perform a 4-fold cross-validation procedure. In this procedure, 3/4 of the data is used to train an MVPA classifier (Fig. 1E, second-to-left panel). The remaining 1/4 of the NRMs are used to determine classifier performance. We performed this computation on a millisecond-by-millisecond basis, evaluating the accuracy of classifier performance as a function of time (Fig. 1E, second-to-right panel). The resulting time courses of decoding accuracy for each laminar compartment were then compared to a randomized trial shuffle control to determine statistical significance (Fig. 1E, rightmost panel).

Before investigating each stimulus dimension in isolation, we evaluated in how far the average laminar profile of spiking activation to our stimuli matched predictions from the CCM (Fig. 2A). To do so, we spatially aligned the spiking data from each recording session to the layer 4C/5 boundary (see Methods and (42)). Using these layer-aligned datasets, we computed the grand average spiking response to all stimuli as a function of cortical depth and time. The resulting laminar profile of activation was consistent with both the expectations set by the CCM and previous studies of laminar visual activation (12, 27, 43–45). This pattern of sensory activation occurs regardless of stimulus dimension, raising the question of how stimulus-specific information is extracted within this activation sequence. To answer this question, we applied MVPA using a “moving searchlight” analysis (46). Specifically, we limited both our training and test data sets to three neighboring electrode channels, performed MVPA over time, and then repeated the process after moving this “searchlight” 100 microns deeper along the electrode array. In this analysis a classifier is trained and tested for each timepoint of the response, in 1 ms increments. (see Methods). The results of this approach are shown in Fig. 2C.

**Figure 2:**
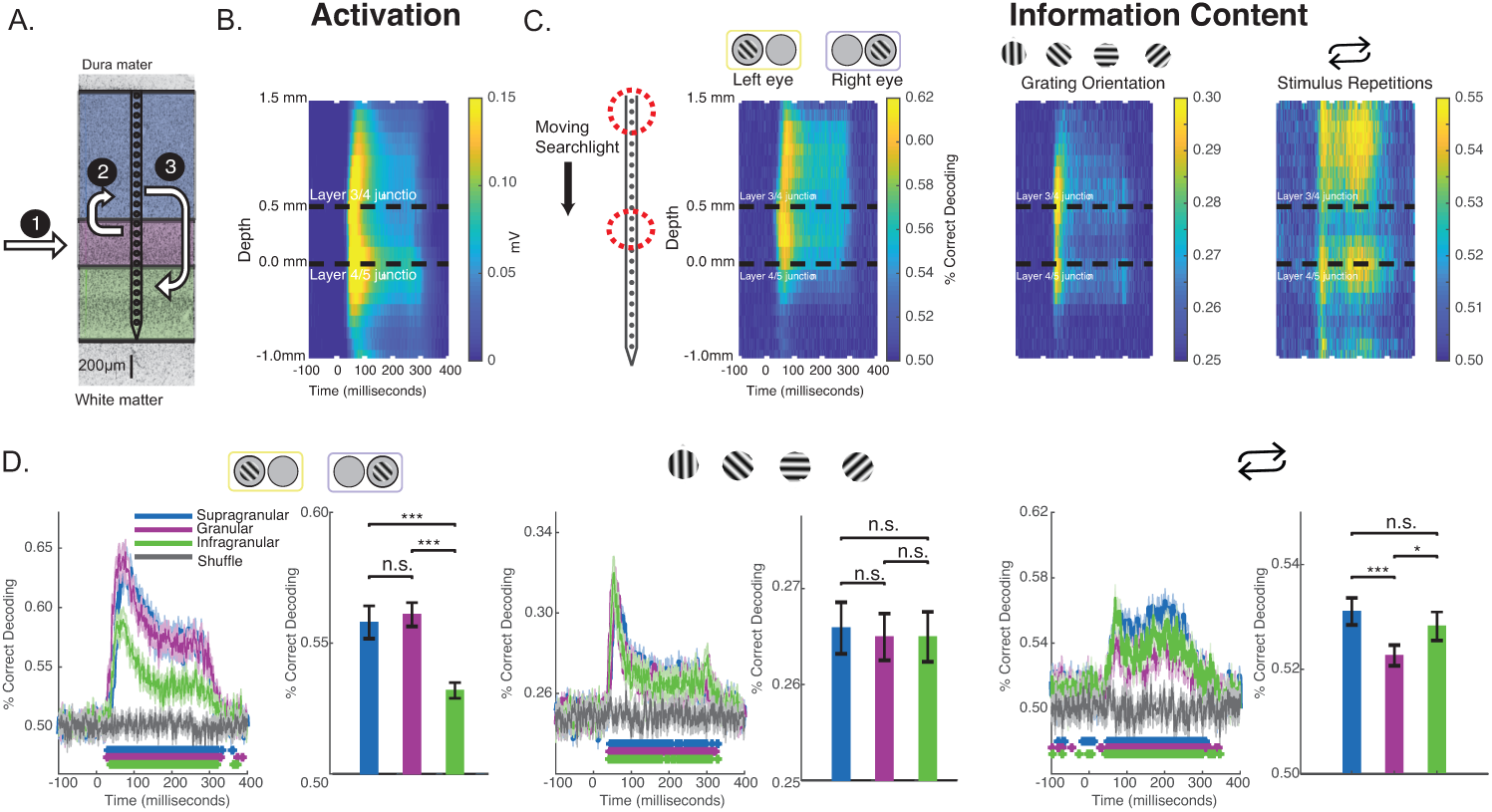
(A) Canonical microcircuit model (CCM) of neural activation in V1. Feedforward activation initially excites the middle layers before reaching upper and lower layers of cortex. (B) Grand average laminar MUA profile to all stimulus presentations along the depth of the electrode (all sessions, both monkeys). (C) Decoding performance using a moving searchlight along the electrode array for eye of origin (leftmost panel), grating orientation (middle panel), and stimulus repetition (rightmost panel). (D) Time series of MVPA decoding for eye of origin (leftmost panel), grating orientation (middle panel), and stimulus repetitions (rightmost panel). Graphs show decoding accuracy as a function of time and laminar compartment, together with a randomized shuffled control as a baseline. Significance is indicated with colored asterisks above the abscissa using Wilcoxon signed-rank test, FDR corrected, q<0.01. Bar plots to the right indicate time-averaged statistics of the data with Wilcoxon signed-rank test P values (* p<0.05, ** p<0.01, *** p<0.001) above the plots.

We first focused on the eye-of-origin for each stimulus presentation. While V1 harbors both neurons that respond to one or both eyes, most of the neurons that respond to one eye only (monocular neurons) are located in the middle, granular layers (7). This finding is consistent with neuroanatomy, as the granular layers receive the bulk of (monocular, eye-specific) inputs from the lateral geniculate nucleus of the thalamus (LGN) that connects eye and cortex (47). As outlined in the Introduction, a long-standing, popular hypothesis is that the eye-specific inputs in the middle layers are merged to a combined (binocular) response in the layers above, even though most V1 neurons maintain preference for one eye over the other (15, 48–50). Neurons in the uppermost layers of V1 project to neurons in V1’s lower layers, so if the upper layers form a combined binocular signal, this signal should be present in the lower layers as well (15, 51, 52). However, based on several other pieces of empirical evidence, an alternative hypothesis postulates that the two eyes’ signals are merged at or before LGN responses arrive in the middle layers of V1 (for review see (53)).

Using MVPA to adjudicate between these hypotheses, we found that the information regarding eye-of-origin initially followed the CCM profile of general activation, with neurons reliably indicating whether a stimulus was shown to left or right eye in the middle layers, followed by the upper layers of V1. Surprisingly, this eye-specific information largely diminished once neuronal activation reached the lower layers of V1 (Fig. 2C, left panel). These timing differences can clearly be seen for a layer-specific MVPA using all electrode channels within the middle, upper and lower layers of V1, respectively (Fig. 2D). We utilized this analysis to perform several statistical comparisons. First, we compared decoding performance on a millisecond-by-millisecond basis against a randomized trial shuffle control (see colored asterisks below time series). Second, we compared decoding across laminar compartments (see bar graph to the right). Decoding of eye-of-origin first emerged in the middle layers (29ms), followed by the upper (40ms) and lower layers (40ms). Decoding which eye the stimuli were shown to was comparable between middle and upper layers but significantly reduced in the lower layers, suggesting that eye-specific information is largely preserved when granular neurons project to neurons in the layers above. However, decoding of eye-of-origin is relatively poor in the lower layers of V1, suggesting that, at least on the multiunit-level, there is significant binocular convergence after activation reaches the upper layers of cortex. This finding is consistent with the original hypothesis that merging of eye-specific signals predominantly occurs in the upper layers of V1, which then project a largely combined binocular signal to the lower layers.

Next, we computed the laminar evolution of stimulus orientation information. As outlined in the Introduction, one of the original hypotheses regarding the functional layout of V1 states that orientation selectivity (tuning) is less pronounced in the middle layers of V1 (7, 15). Yet, several authors have challenged this idea, arguing that V1 already receives orientation-tuned inputs (21–26). We thus wondered what the laminar profile of MVPA-based decoding of stimulus orientation across V1 layers might be, especially since this question cannot be fully answered by contrasting two stimulus conditions using standard univariate techniques.

We binned our grating stimuli into four groups (0 degrees, 45 degrees, 90 degrees and 135 degrees, respectively) and trained a classifier to discriminate between them. Classifier performance as a function of cortical depth and time is shown in the middle panel of Fig. 2C. Interestingly, we found that information regarding stimulus orientation was more transient than information regarding of eye-of-origin. Moreover, the laminar profile was strikingly different: the center of the granular layers discriminated relatively poorly between gratings of varying orientation, and neurons in the layers above and below did so without any significant temporal delay. Closer inspection of the layer-resolved decoding (middle panel of Fig. 2D), collapsed across time, revealed that there was no significant difference between any of the laminar compartments (bar plots). These results seem to suggest that stimulus orientation information is extracted almost uniformly across V1 layers. However, visual inspection reveals clear differentiation within the middle layers, which is lost when collapsing this layer into a single measure. This heterogeneous pattern within the granular layers might at least be partially explained by the fact that the middle layers host several sublayers that each receive separate inputs from the LGN (47), although it is not immediately clear how the granular sublayers relate to the specific pattern we found.

Given that V1 is known to modulate its responses depending on contextual cues, such as the behavioral state of the animal or stimulus history (12, 14, 41), we next examined how stimulus history affects the laminar flow of stimulus-specific information. To do so, we first studied the laminar flow of information of whether a stimulus was novel or preceded by another stimulus in the stimulation sequence. We found that this information regarding stimulus history yielded yet another pattern of laminar information flow (Fig. 2C, rightmost panel). We found that the bulk of information regarding stimulus history resided outside the granular input layers. This finding was also apparent in layer-specific MVPA (Fig.2D, rightmost panel). These results are in line with earlier work showing that V1 granular layers are least affected by the adaptive effects of repeated visual stimulation (41). However, our current results go beyond this observation by showing that the information regarding stimulus history did not exhibit the typical decay following the initial transient, potentially suggesting involvement of some kind of recurrent process.

To further investigate this potentially recurrent process, we decoded neuronal data based on a classifier that was trained for another time period of the same neuronal response (“time generalization”) (54, 55). The result of this analysis is a 2D “time generalization matrix” that plots training time against decoding time. Fig. 3A, illustrates several possible outcomes for generalization matrices. It is possible, for example, that there is little to no generalization between a classifier trained at one time and tested on the remaining time of a neuronal response. In other words, spiking might be constantly changing in a way that any information used to discriminate between stimuli is specific to each individual point in time of the neuronal response (“unique states”). In contrast, if the information used to discriminate between stimuli were static across the neuronal response, we would expect a square-like pattern (“sustained”). This analysis can also show information decaying over time (“information decay”). An asymmetric pattern occurs because a classifier trained on lower SNR data generalizes better to higher SNR data than the converse (56). Lastly, information might reoccur at a later time point of a response (“recurrence”).

**Figure 3:**
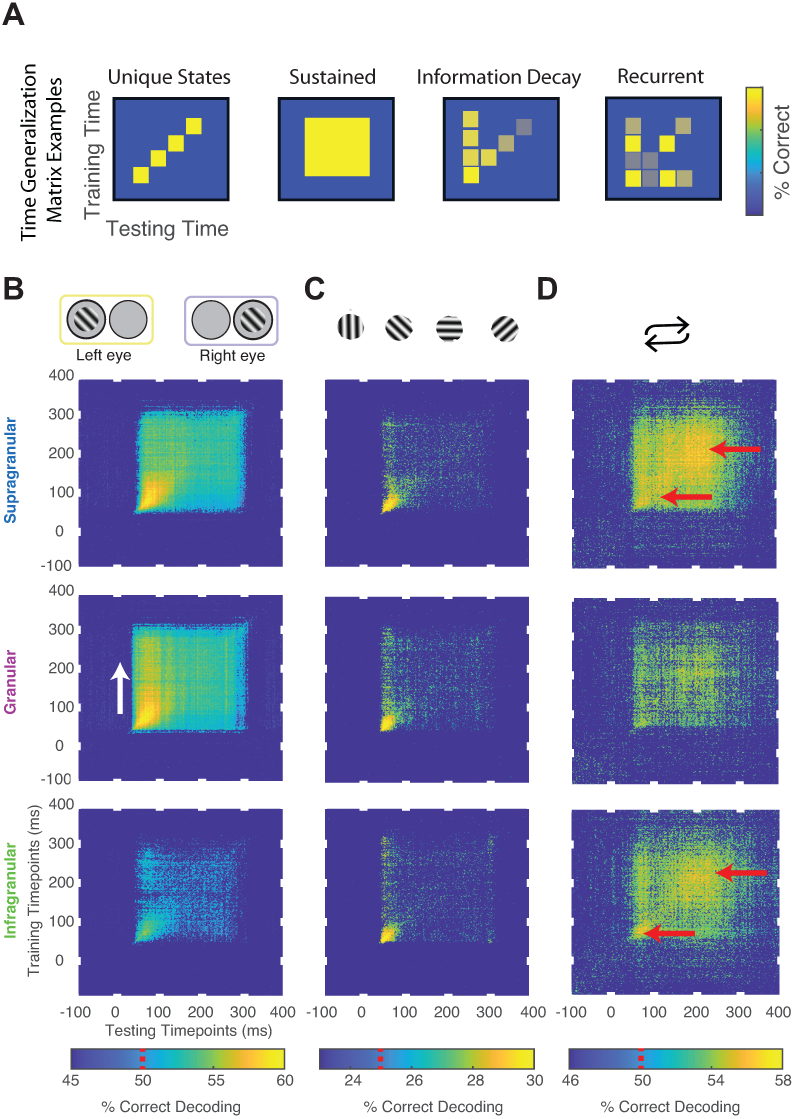
Time generalization results. (A) Cartoon models of possible results. (B) Time generalization analysis performed for: (B) Eye-of-origin, (C) Orientation Grating, (D) Stimulus repetitions (see Methods for details). Chance decoding level is indicated on each color bar by a red line. Red and white arrows are added for emphasis.

We performed time generalization analysis for the decoding of eye-of-origin, stimulus orientation as well as stimulation history within each laminar compartment (Fig. 3 and Suppl. Fig. 2). Decoding eye-of-origin (Fig. 3, left column) was mostly sustained but also exhibited some information decay within each laminar compartment (Fig. 3B, leftmost columns). Decoding of stimulus orientation, in contrast, was less sustained (Fig. 3, middle column). Interestingly, whether or not a stimulus preceded or succeeded other stimuli showed a very different pattern. Specifically, the time generalization matrix was suggestive of recurrent processing, in that the initial information emerges, weakens and then re-emerges at a later time point. This reactivation pattern was most prominent in the supragranular and infragranular layers (Fig. 3, rightmost column). This findings confirms the notion that information regarding stimulus history reverberates in the feedback-recipient layers of V1.

Given that information regarding stimulus history reverberates within the CCM in a layer-specific way, we next studied how stimulus history might *interact* with the layer-specific information processing for stimulus orientation and eye-of-origin. Specifically, we tested how well neuronal responses in each laminar compartment discriminated between stimulus orientation and eye-of-origin, depending on whether the stimuli were first in the sequence or appeared as a repetition. The result is shown in Fig. 4. Intriguingly, classifier performance for stimulus orientation significantly increased when a stimulus was a repeat. This finding is in line with the finding that cortical adaptation increases sharpening of orientation tuning curves (57). Surprisingly, though, we found that this is not the case for eye-of-origin information. Indeed, discriminability between the eyes significantly decreased for repeated stimuli (Fig. 4). This shows an intriguing differentiation for the effects of visual adaptation, in that neurons across all layers of V1 get more sensitive for the orientation of visual stimuli but less sensitive to which eye these stimuli are presented.

**Figure 4:**
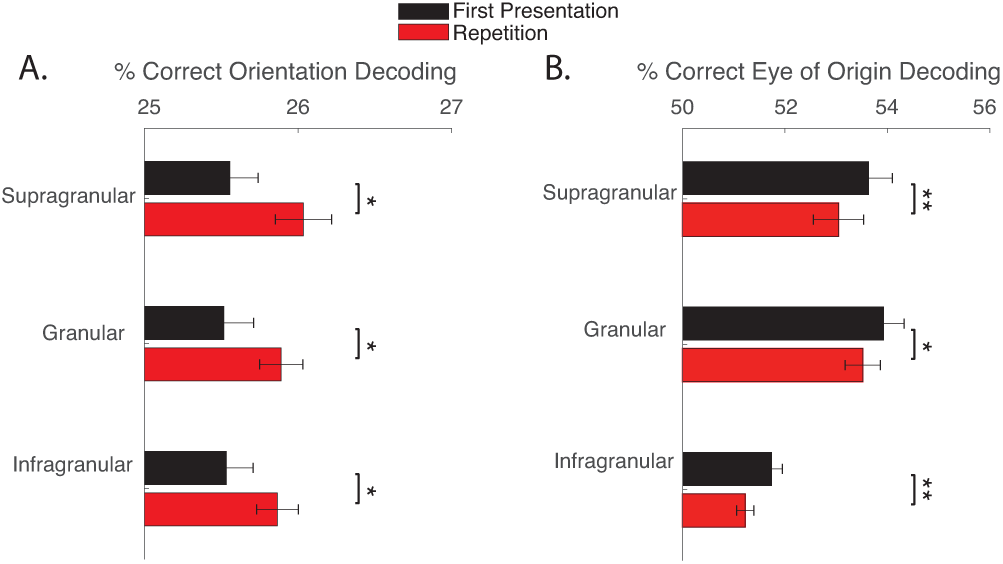
Decoding of (A) Orientation and (B) Eye-of-Origin for initial presentations (black) and repetitions (red) as a function of laminar compartment (ordinate), averaged over the entire response period (400ms following stimulus onset). Wilcoxon signed-rank test P values (* p<0.05, ** p<0.01, *** p<0.001) above the plots.

## Discussion

Recent studies using linear multielectrode arrays in V1 have successfully contrasted externally evoked feedforward activation with internally generated feedback (11–14, 44). These results are encouraging as they demonstrate that the flow of neural activation across cortical layers is highly informative regarding the context of neuronal activation – an important insight that is largely absent in single electrode recordings. In this study we went significantly beyond these earlier findings by showing how the dynamic build-up of cortical laminar activation contains several parallel streams of information about specific stimulus dimensions that are difficult to trace using univariate analyses, even when laminar data has been obtained.

Our findings are in line with the hypothesis that each eye’s stream of information stays largely separate until visual activation reaches the upper layers of V1. We found a drastic reduction in eye-specific information in the lower layers of V1, suggesting that visual signals are much more strongly binocularly integrated once they reach the deep layers. This pattern is distinct from earlier reports, locating the bulk of V1 binocular neurons in both the upper and lower layers (7). This apparent paradox might be explained by our recent finding that a large fraction of monocular V1 neurons are actually sensitive to both eyes (51), suggesting that a neuron’s preference for one or the other eye is not predictive of how it responds to binocular stimulation (see also (50)).

Furthermore, our results revealed a fine-grained spatiotemporal laminar pattern of orientation tuning, with some but not all sublayers of granular layer 4C exhibiting less sensitivity to stimulus orientation than the superficial and deep layers of V1. Although it is not immediately clear how the specific pattern produced by MVPA relates to the magno- and parvocellular sublayers, our finding seems to be generally in line with the idea that V1 receives at least some LGN inputs that are somewhat “biased” towards certain stimulus orientations, with further processing within V1 producing the more discerning orientation tuning that characterizes this area.

It is also interesting to note that stimulus repetition yielded a unique signature of time generalization in the feedback recipient layers of V1. The temporal features of this pattern are somewhat reminiscent of prior descriptions of feedback modulation in V1 (12). However, our finding goes beyond the demonstration of a secondary peak in activation by revealing that the information content within this activation is specific to contextual information. We also found that information regarding line orientation increases across all layers of V1 when stimuli are shown repeatedly, echoing earlier work that suggested “sharpened” orientation tuning following visual adaptation (41). Yet, information regarding eye-of-origin simultaneously decreased, suggesting that visual adaptation triggers a push-pull like mechanism in V1 where the processing of some stimulus features gets heightened at the cost of others. Interestingly, eye-specific information also seemed to decrease in both the searchlight decoding and time generalization results, indicating that it is more readily dispensed by V1’s CCM compared to other types of stimulus information, which seems in line with the fact that eye-of-origin information is of low behavioral relevance (58). Thus, our results show how information flow across the CCM reflects the functional utility of a cortical area.

## Materials and Methods

### Animal Care and Surgical Procedures

Data from this study were collected from two macaque monkeys (Macaca radiata, one female). A detailed description of the surgical procedures can be found in a previous publication (41). Briefly, in a series of surgeries, each monkey was implanted with a custom MRI-compatible headholder and recording chamber over perifoveal V1 concurrent with a craniotomy. All procedures were in compliance with regulations set forth by the Association for the Assessment and Accreditation of Laboratory Animal Care (AALAC), approved by the Vanderbilt University Institutional Animal Care and Use Committee, and followed National Institutes of Health guidelines.

### Behavioral Paradigm

In each recording session, monkeys were positioned in front of a 20” CRT monitor (Diamond Plus 2020u, Mitsubishi Electric Inc.) operating at 60 or 85Hz. Monkeys passively fixated within a one-degree radius around a central fixation dot and viewed stimuli through a custom mirror stereoscope so that stimuli could be viewed monocularly or binocularly. Stimuli were generated using MonkeyLogic (59, 60) via MATLAB (R2012, R2014a, The Mathworks, Inc.). running on a computer using a Nvidia graphics card. Following 300ms of fixation, monkeys viewed five sequentially presented stimuli for 200ms each, with a 200ms inter-stimulus interval (ISI). If fixation was maintained throughout the five presentation, the monkey was rewarded with juice and relieved of the fixation constraint for an inter-trial interval (ITI). If the monkey broke fixation during trial performance, the presentation was eliminated from analysis and the monkey experienced a short timeout (1-5s) before starting the next trial. Each stimulus in the presentation sequence was a sinusoidal bar grating of equivalent size, spatial frequency, and phase, with variable orientation. For each recording session, the stimuli were optimized for the measured neural activity evaluated by listening to the multi-unit activity (MUA) during exposure to a wide variety of stimuli. We selected stimulus parameters that evoked the greatest neural response. For a more detailed description of the paradigm, as well as further information on stimulus optimization and receptive field mapping, see previous publications (14, 41, 61).

### Neurophysiological Procedure

All data used in this paper are available upon request from the communicating author, pending approval by Vanderbilt University. During task performance, broadband (0.5Hz-12.207kHz) intracranial voltage measurements were taken at a sampling rate of 30kHz and amplified, filtered, digitized using a 128-channel Cerebus™ Neural Signal Processing System (NSP, 409 Blackrock Microsystems LLC). Neuronal data was downsampled offline to 1kHz, following low-pass filtering with an anti-aliasing filter. Gaze position was recorded at 1kHz (NIDAQ PCI-6229, National Instruments) using an infrared light sensitive camera and commercially available eye tracking software (Eye Link II, SR Research Ltd.; iView, SensoMotoric Instruments). Recordings took place inside an electromagnetic radio frequency-shielded booth and were performed using one or two acute laminar multielectrode arrays with 24 or 32 contacts with 0.1mm electrode spacing and impedances ranging between 0.2 and 0.8MOhm at 1kHz (U-Probe, Plexon, Inc.; Vector Array™, NeuroNexus). Electrodes were connected to the NSP using analog headstages. In each recording, the electrode array(s) were introduced into dorsal V1 through the intact dura mater using a chamber-mounted microdrive (custom modification of a Narishige International Inc. Micromanipulator) and adjusted such that the majority of recording contacts spanned the cortical sheet.

### Laminar Alignment

CSD in response to brief visual stimulation was used to find the boundary between the granular and infragranular compartments of V1 as per previously documented methods (11, 14, 27, 41, 45, 62). Additional neurophysiological criteria were used, such as well-defined patterns of LFP power spectral density (12, 41, 63), signal correlations between LFP recorded on differing channels, and latency (64) of stimulus-evoked MUA. The granular to supragranular boundary was set to 0.5mm above the granular to infragranular boundary.

### Preprocessing

All recording channels found to be within the brain were taken and multiunit signals were computed. Channels in the brain were found by determining first whether a visual response could be evoked on the channel and second, whether a receptive field was present for the multiunit and/or LFP activity through a previously described receptive field mapping paradigm (41). If the channel was found to be in the brain, the broadband neural signal recorded at that channel was then band-pass filtered between 500 and 5000Hz, rectified, and low-pass filtered at 200Hz using Butterworth filters (64, 65). These derived neural signals were then used in performing both the univariate and multivariate analyses.

### Multivariate Pattern Analysis

Following data preprocessing, we used CoSMoMVPA (66) to analyze the data with a time-resolved decoding approach for eye of origin, orientation, and stimulus history (repetitions). A classifier was trained and tested along each time point using linear discriminant analysis for each of these distinctions. We used 4-fold cross validation with a ratio of three (training) to one (test) to evaluate decoding accuracy. In this procedure, trials were randomly assigned to one of four subsets of data. Three of the four subsets were then pooled together to train the classifier. Then, decoding accuracy was tested on the remaining subset. This procedure was repeated four times, such that each of the subsets was tested at least once. Decoding results are reported in percent correct classifications at each time point of the time series [-100ms to 400ms]. This analysis was conducted independently for each record460 ing session (n=48), stimulus dimension of interest (eye of origin, orientation, and repetitions), and layer (supragranular, granular,and infragranular). Mean and standard error were then calculated across recording sessions at each time point. Each time point was tested for significant above-chance decoding using a Wilcoxon signed rank test. To correct for multiple comparisons, we used the false discovery rate adjusted p-values with alpha=.01. Chance decoding was calculated by using a shuffle control, in which the labels for each stimulus presentation is randomized. For each of the decoding distinctions, the subsets were balanced, such that both training subsets and testing subsets contained the same number of trials for each stimulus category. For orientation decoding, all recording sessions were used for analysis. However, some recording sessions included orientation presentations that were not shown in other recording sessions (i.e. 22.5 degrees in one recording session and 30 degrees in another sessions).

Therefore, orientation presentations were binned into four categories: 0-44 degrees, 45-89 degrees, 90-134 degrees, and 135-179 degrees. For trial repetition decoding, the five stimuli presentations for a given trial were grouped as either the first presentation or as a repetition. To have an equal number of first presentations and repetitions, we randomly subsampled from the repetitions to match the number of first presentations. For the time generalization analysis, a similar decoding analysis was performed as described above for each of the stimulus dimensions. However, the classifier trained was performed on one time point for each stimulus dimensions and tested on each of the remaining timepoints. This procedure was repeated until all timepoints were used as training timepoints. To determine the effects of repeated stimuli presentations on orientation and eye of origin decoding, we further divided the repetition subset of data into balanced eye of origin subsets and balanced orientation subsets. We then again performed a 4-fold classification using a linear discriminant analysis classifier.

## ACKNOWLEDGMENTS

A research grant from the National Eye Institute (1R01EY027402-02) supported this study. The authors would like to thank B. and R. Williams, M. Schall, and M. R. Feurtado for technical support. D.A.T. is supported by a by NIGMS of the National Institutes of Health (T32GM007347). We thank L. Daumail and B. Mitchell for comments on an earlier version of this manuscript. J.A.W. and K.D. are supported by a National Eye Institute Training Grant (5T32EY007135-23).

## Author’s Contributions

D.A.T., J.A.W., and A.M. conceptualized the study. M.T.W., T.C. and A.M. supervised the study. J.A.W., M.A.C. and K.D. collected the data. J.A.W., M.A.C. and K.D. preprocessed the data. D.A.T. performed the main data analyses and created all figures. D.A.T., J.A.W. and A.M. wrote the initial draft of the paper. D.A.T., J.A.W., M.A.C., K.D., T.C., M.T.W. and A.M. revised the paper and approved the final version of the manuscript.

## Supplemental Figures

**Supplemental Figure. 1:**
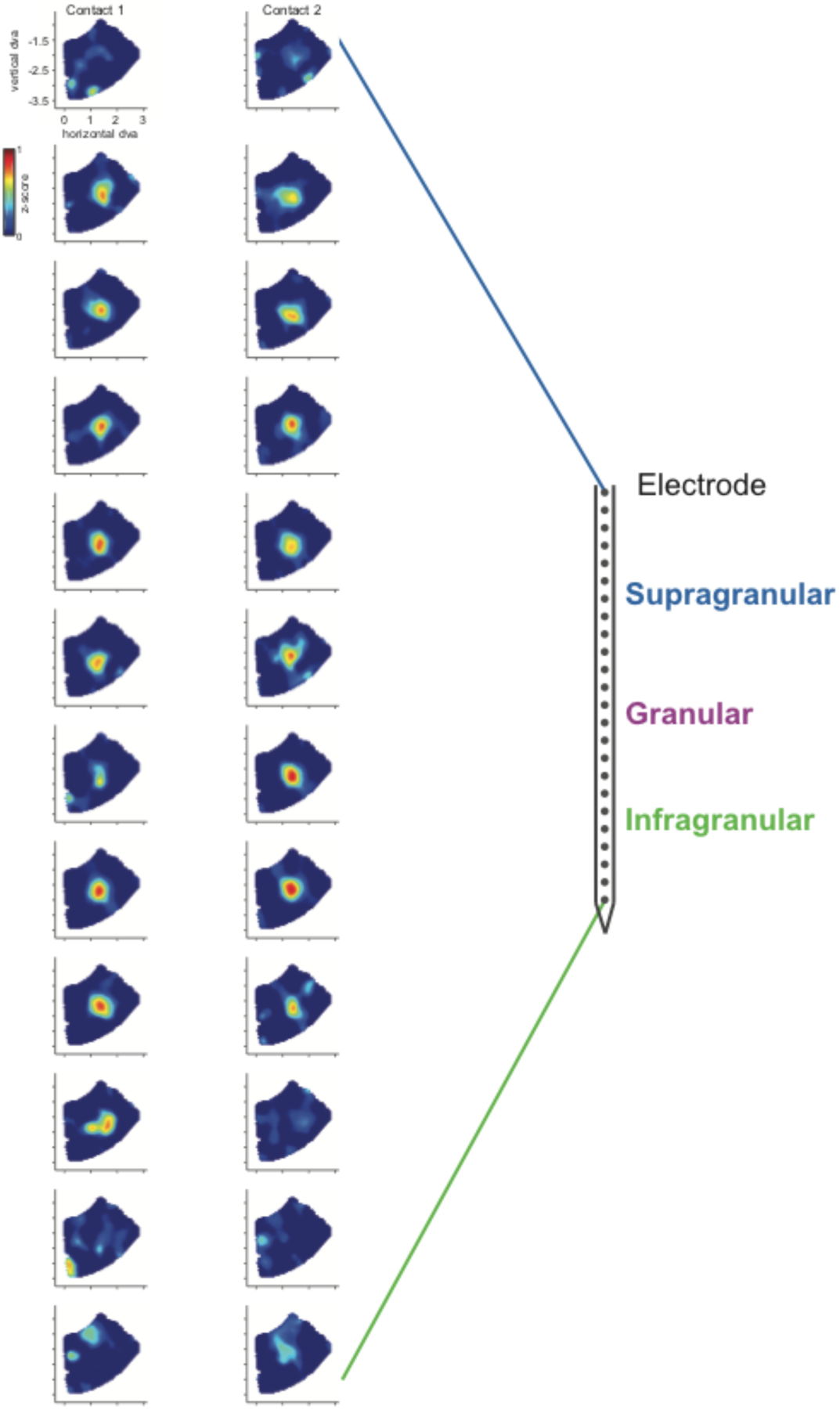
Receptive Field Mapping. For each contact of the linear multielectrode array, we computed the magnitude of MUA spiking responses as a function of visual field stimulation using a reverse-correlation technique (see Methods). Colored plots to the left show averaged neuronal response in units of standard deviation as a function of angle and magnitude in visual degrees. Panels are arranged in descending order with each column representing neighboring channels on the electrode array so that each row represents the electrode channel that is 200 microns below the channel above. Note that the receptive field locations deviate little between the top and the bottom of the array, indicating that the electrode was inserted perpendicularly to the cortical surface. The topmost and bottommost channels of the array produced no visual responses as these electrode channels reached outside the cortical thickness.

**Supplemental Figure 2:**
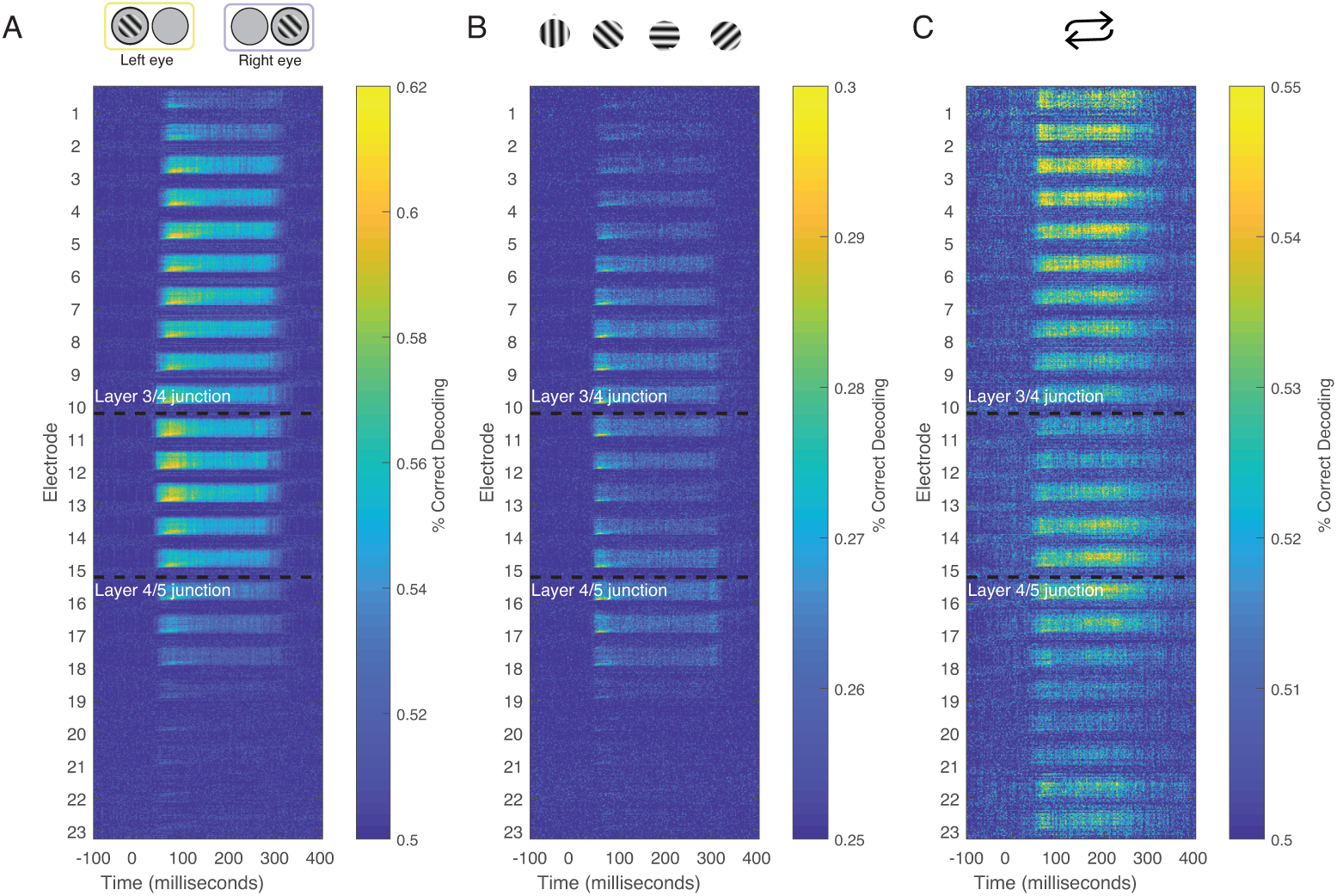
Combined time generalization and moving searchlight analysis along the depth of the linear electrode array. We performed this analysis for each of the main stimulus dimensions analyzed in this paper: Stimulus (A) eye-of-origin, (B) orientation and (C) repetition. Each sub-panel shows a series of time generalization plots ranging from 100ms before stimulus to presentation to 400ms post stimulus presentation using a moving searchlight of three electrodes and two electrodes at the end of the electrode array.

